# Sex-Related Gene Expression in the Posterodorsal Medial Amygdala of Cycling Female Rats Along with Prolactin Modulation of Lordosis Behavior

**DOI:** 10.1101/2024.11.12.623226

**Authors:** Letícia Bühler, Ana Carolina de Moura, Márcia Giovenardi, Vincent Goffin, Alberto A. Rasia-Filho

## Abstract

The rat posterodorsal medial amygdala (MePD) is sexually dimorphic, has a high concentration of receptors for gonadal hormones and prolactin (PRL), and modulates reproduction. To unravel genetic and functional data for this relevant node of the social behavior network, we studied the expression of *ERα, ERβ, GPER1, Kiss1, Kiss1R, PRGR, PRL, PRLR, EGR1, JAK2, STAT5A,* and *STAT5B* in the MePD of males and females along the estrous cycle using the RT-qPCR technique. We also investigated whether PRL in the MePD would affect the sexual behavior display of proestrus females by microinjecting saline, the PRL receptor antagonist Del1-9-G129R-hPRL (1 µM and 10 µM), or PRL (1 nM) and Del1-9-G129R-hPRL (10 µM) 3h before the onset of the dark-cycle period. The estrogen-dependent lordosis behavior, indicative of sexual receptivity of proestrus females, was recorded and compared before (control) and after (test) microinjections in these groups. Sex differences were found in the right and left MePD gene expression. *ERα* and *Kiss1R,* as well as *PRL, Short PRLR*, and *STAT5B* expression, is higher in cycling females than males. *Kiss1* expression is higher in males than females, and *GPER1* is higher during diestrus than proestrus. Furthermore, Del1-9-G129R-hPRL in the MePD significantly reduced the full display and quotient of lordosis in proestrus females, an effect restored by the co-microinjection of PRL. In conjunction, the expression of studied genes showed specific sex and estrous cycle phase features while, in proestrus, PRL action in the MePD plays an essential role in the display of lordosis during the ovulatory period.

**Highlights:**

- The MePD expression of *Kiss1* is higher in males than females.
- *ERα* and *Kiss1R* expression is higher in cycling females than males.
- *PRL, Short PRLR,* and *STAT5B* expression is higher in cycling females than males.
- *GPER1* expression is higher during diestrus than in proestrus.
- Del1-9-G129R-hPRL action in the MePD reduced lordosis quotient during proestrus.

## 1. Introduction

The rat posterodorsal subdivision of the medial amygdaloid nucleus (MePD), lateral to the optic tract and ventral to the stria terminalis, is part of the “extended amygdala” and a social behavior network that modulates social behaviors in males and females [1–3,11,16]. Local cells are notably responsive to circulating levels of androgens and display estradiol α and β receptors (ERα, ERβ) and progesterone receptors (PRGR) [1,2,4]. The MePD neurons and astrocytes are sexually dimorphic and show morphological (in dendritic spine density), epigenetic, and electrophysiological plasticity during the estrous cycle [1,5–9]. This forebrain area processes chemosensory olfactory and vomeronasal inputs [2,10], alters aggressive display [11], and evokes sympathetic and parasympathetic reflex responses [12] and neuroendocrine responses to stressful stimuli [13]. It also modulates the occurrence of puberty [14] and sexual (in both males and females) [1–3,11,15], and maternal behavior [5,16] by integrating activity with various hypothalamic nuclei [17].

In males, the MePD participates in penile erection, intromission, and ejaculation [11,18,19]. In females, the hormonally-mediated cyclically switches in the MePD reduce the number of dendritic spines, alter the excitatory and inhibitory inputs, the output activity, and likely disinhibits female sexual behavior, integrating the occurrence of lordosis behavior during mating with the ongoing, timely hypothalamic gonadotropin-releasing hormone (GnRH) secretion for ovulation [1,5,8,11,16,19]. In this regard, plasma levels of follicle-stimulating hormone (FSH) and luteinizing hormone (LH) change during the estrous cycle phases, peaking at proestrus. Likewise, estradiol levels gradually increase during diestrus, reach maximum levels during proestrus, and return to baseline during estrus; progesterone peaks during proestrus after the estradiol surge and returns to lower values afterward; and prolactin (PRL) has the first and highest peak during proestrus and a second and lower one during estrus [20,21].

Gonadal hormones regulate gene transcription in the nervous tissue by acting on their receptors and inducing complex, integrated intracellular signaling pathways. For example, estradiol can synergistically induce the expression of PRGR and alter brain synaptic plasticity [22,23]. The transcription factor STAT5 (signal transducer and activator of transcription 5) is related to ERα actions [24] as well as is activated by PRL via Janus kinase 2 (JAK2) [25], whose inhibition has been shown to impair the ability of PRL to increase *ERα* expression [26]. The G protein-coupled estrogen receptor (GPER1) is part of the estradiol-mediated effects on spinogenesis, female sexual behavior, and social learning [27]. GPER1, although acting independently of ERα and ERβ, activates common second messenger pathways to exert genomic effects [28]. Furthermore, a specific form of the neuropeptide kisspeptin, kisspeptin 1 (KISS1), is encoded by the *Kiss1* gene [29]. This neuropeptide signaling modulates the hypothalamic-pituitary-gonadal axis and the occurrence of sexual behavior. In the medial amygdala (MeA) of mice and rats, *Kiss1* gene expression is sexually dimorphic, varies with the estrous cycle, is highly regulated by sex steroids, and involves ERα activity [29–32]. The KISS1 ligand is the kisspeptin 1 receptor (KISS1R), which shows abundant expression in the amygdala of rats [33]. However, additional data on gene expression involved with male-to-female differences and ovarian steroids’ cyclic actions, specifically in the MePD, are needed.

Interestingly, the rat MePD also has a high density of PRL receptors (PRLR) [34] and immunoreactive PRL fiber projections within the MeA neuropil [35], but the functional role of PRL in the MePD is not known. PRL is fundamental to lactation but also acts as a neuromodulator, affecting synaptic plasticity and promoting neurogenesis and neuronal cell survival [36]. PRL can be synthesized and secreted by extrapituitary tissues [37,38] to bind to membrane receptors with short or long cytoplasmic domains [39] and possibly different signaling pathways activated by each isoform [40]. Estrogen regulates PRLR expression [41], and the effects of PRL involve the induction of mitogen-activated protein kinase (MAP kinase)-dependent Early Growth Response Factor 1 (EGR1) [42]. EGR1 expression has sex differences and variations across the estrous cycle [43] relevant to brain plasticity [44]. PRL is also involved in specific increases in tyrosine phosphorylation of JAK2 and subsequent phosphorylation of STAT5B, but not STAT5A, in neurons [25]. Furthermore, acute intracerebral administration of PRL stimulates estrogen-dependent lordosis behavior, indicative of female sexual receptivity [45]. In contrast, the administration of bromocriptine, a dopamine D2 receptor agonist that inhibits PRL secretion during proestrus, suppresses it [46]. Based on these data, we tested whether or not PRL acting in the female MePD modulates the display of lordosis behavior along with the highest levels of sex hormones and PRL in circulation during proestrus.

Here, we studied and compared the gene expression of estradiol receptors (*ERα, ERβ,* and *GPER1*); *Kiss1* and *Kiss1R*; *PRGR; PRL; PRLR short and long isoforms; EGR1, JAK2, STAT5A,* and *STAT5B* in the MePD of males and cycling female (in diestrus, proestrus, and estrus). Moreover, we investigate whether PRL action in the MePD would affect the sexually receptive reflex display of lordosis behavior in female rats during proestrus by microinjecting locally a selective antagonist of the PRLR (Del1-9-G129R-hPRL) [47] alone or combined with PRL.

## 2. Material and methods

### 2.1 Animals

Adult Wistar rats (2-3 months old, N = 29 males and 87 females) were obtained from the Federal University of Health Sciences of Proto Alegre (UFCSPA, Brazil) animal facility. Animals of each sex were housed in groups (3-4 animals per cage) with water and food *ad libitum* and under standard laboratory conditions, with controlled room temperature (23 ± 1°C) and 12/12-h light/dark cycle (lights on at 7 a.m.).

All experiments were performed according to international laws regulating laboratory animals’ care. The local Ethics Committee on the Use of Animals/UFCSPA reviewed and approved this study (#616/19 and 641/19),

### 2.2 Gene expression study

#### 2.2.1 Experimental groups and MePD sampling

Animals were divided into four groups: (1) males and (2-4) females in different phases of the estrous cycle (diestrus, proestrus, and estrus). Vaginal cytology was performed daily to determine the regularity of the estrous cycle over two weeks. Only normally cycling females were included in this study.

At the end of this initial period, females were again studied for their estrous cycle phases to be randomly included in one of the experimental groups. In the morning (at 8 a.m.), males and females received an intraperitoneal injection of 240 mg/kg ketamine and 30 mg/kg xylazine and, after being deeply anesthetized, were euthanized by decapitation. The brains were immediately removed from the skull and sectioned. From a single thick coronal slice, the MePD was identified according to the rat brain atlas [48] and removed along a maximum *postmortem* period of 2 min, as previously done [7,49]. Right and left hemisphere samples were studied separately. Due to the reduced size of the MePD, it was necessary to pool tissue from 3 animals to compose each one of a total of 6-8 samples per experimental group. All samples were immediately placed in Eppendorf tubes containing 50 µL of RNAlater® Stabilization Solution (Invitrogen, USA) and refrigerated at 4°C for no more than 24 h. The excess of RNAlater was removed the following day, and the samples were stored at −80°C until the molecular analysis.

#### 2.2.2 Total RNA extraction and RT-PCR

The total RNA from each MePD sample pool was extracted using PureLink® RNA Mini Kit (Invitrogen, USA) according to the manufacturer’s instructions. The total RNA quantification of the samples was performed by spectrophotometry BioSpec-nano® (Shimazu, Japan) at 260nm and 280nm with an optical aperture of 0.7 mm.

After total RNA extraction, complementary DNA (cDNA) synthesis was performed by reverse transcription reactions (GoScript™ Reverse Transcription Kit; Promega, Brazil). A total of 12 µg of RNA was incubated with oligo (dT), dNTPs, and DEPC water (diethyl pyrocarbonate) for 5 minutes at 65 °C, followed by 1 minute on ice. Buffer solution, DTT (dithiothreitol), and RNaseOUT were added for 2 minutes at 37 °C. Finally, the enzyme M-MLV-RT was added for cDNA synthesis for 1 hour at 50 °C. The synthesis reaction was inactivated by incubation at 70 °C for 15 min.

#### 2.2.3 qPCR analysis of gene expression

All primer sequences used in this study were designed using the software Primer-3 based on rat mRNA sequences in the GenBank database. The specificity of the primers was checked using BLAST search against nucleotide collection (nr) of the NCBI database. All primers were purchased from Invitrogen (Brazil) as follows:

***β-Actin***: F: 5’TATGCCAACACAGTGCTGTCTGG’3;
***β-Actin***: R: 5’TACTCCTGCTTGCTGATCCACAT’3;
***ERα***: F: 5’GATGGTCAGTGCCTTATTGGATGC’3;
***ERα***: R: 5’GCAGGTTCATCATGCGGAATCG’3;
***ERβ***: F: 5’CAATCATCGCTCCTCTATGC’3;
***ERβ***: R: 5’GGCCTTACATCCTTCACATGA’3;
***GPER1***: F: 5’TGCAAGCAGTCTTTCCGTCA’3;
***GPER1***: R: 5’CTCGTCTTCTGCGCCACATA’3;
***Kiss1***: F: 5’TGCTGCTTCTCCTCTGTGTGG’3;
***Kiss1***: R: 5’ATTAACGAGTTCCTGGGGTCC’3;
***Kiss1R***: F: 5’CTTTCCTTCTGTGCTGCGTA’3;
***Kiss1R***: R: 5’CCTGCTGGATGTAGTTGACG’3;
***PRGR***: F: 5’CGATGGAAGGGCAGCATAAC’3;
***PRGR***: R: 5’CGAGGGCTCTCATAACTCGG’3;
***PRL***: F: 5’TTCTTGGGGAAGTGTGGTC’3;
***PRL***: R: 5’TCATCAGCAGGAGGAGTGTC’3;
***PRLR (both isoforms)***: F: 5’CTGGGCAGTGGCTTTGAAG’3;
***Short PRLR***: R: 5’AAGGGCCAGGTACAGATCCA’3;
***Long PRLR***: R: 5’CCAAGGCACTCAGCAGCTCT’3;
***EGR1***: F: 5’GAGCACCTGACCACAGAGTC’3;
***EGR1***: R: 5’AAAGGGGTTCAGGCCACAAA’3;
***JAK2***: F: 5’TATGCTCCCGAATCCTTGAC’3;
***JAK2***: R: 5’TCTGCCCTTGTTTATCATTGC’3;
***STAT5A***: F: 5’GCCCTCAGGCTCACTACAAC’3;
***STAT5A***: R: 5’AAAGGCGGGGGTCAAGACT’3;
***STAT5B***: F: 5’TGGGGCAGAACGAGGTTGTA’3;
***STAT5B***: R: 5’ACCTCGATGGGGAAATGCTG’3.

The analysis of the expression of the target genes was performed at the transcriptional level, in which the mRNA content was evaluated, using specific primers to amplify the cDNAs (10 ng of cDNA per well) of the genes studied. Actin beta (ActB) was used as a housekeeping gene for internal control of gene expression [reference genes in accordance to 50]. Amplification products were analyzed using the SYBR™ Green method by Real-Time PCR (Life Technologies, Brazil). The Ct value of each reaction was used to calculate the level of mRNA expression of a specific gene after normalization compared to the expression of the reference gene that was analyzed in parallel on the same reaction plate. The relative quantification (mean of normalized expression to the housekeeping gene) was calculated by the 2^-ΔΔCT^ method [51].

### 2.3 Female sexual behavior test

#### 2.3.1 Animals and experimental groups

Adult Wistar rats (2-3 months of age, N = 30 sexually experienced males to test female sexual receptivity and 25 females) were obtained from the same federal facility mentioned above. Vaginal cytology was performed daily, and only females showing regular estrous cycles were included in the experimental groups. Females composed four experimental groups. They were microinjected into the right MePD with Vehicle (Veh, 0.9% sterile saline, n = 6); 1 µM of Del1-9-G129R-hPRL (n = 6); 10 µM of Del1-9-G129R-hPRL (n = 7); and co-microinjection of PRL 1 nM and Del1-9-G129R-hPRL 10 µM (n = 6). The volume microinjected was 0.3 µL in all groups. Because there are no available data about the PRL level in the rat MePD, doses were calculated from the amount of PRL circulating during the afternoon of proestrus and the likely diffusion of the PRLR antagonist in the nervous tissue. Both PRL and PRLR antagonists were produced by recombinant technology in bacteria and were chromatography-purified as previously described [47].

#### 2.3.2 Estrous cycle normality and sexual behavior testing

Adult virgin females had their vaginal cytology studied daily (at 12:30 pm) for three consecutive weeks. Accordingly, the estrous cycle phases were classified as diestrus, proestrus, and estrus [20]. The experimental design is shown in Figure 1.

**Figure 1.**
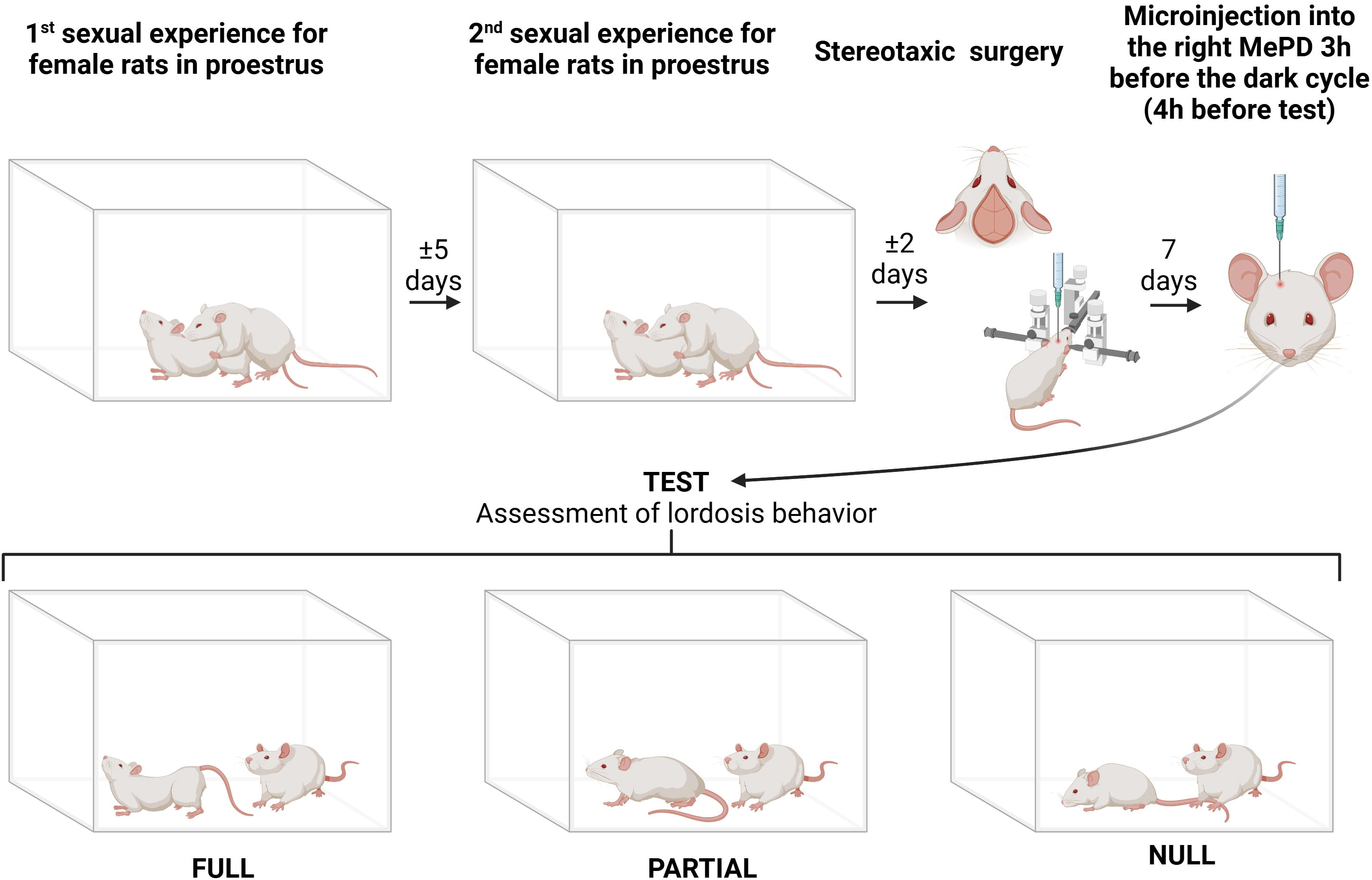
Experimental procedures for testing female sexual behavior. Normally cycling females were submitted to two mating sessions with sexually experienced male rats. Afterward, females were stereotaxically implanted with a cannula directed to the right posterodorsal medial amygdala (MePD). Following recovery, females were tested again for sexual receptivity in proestrus. Females would show no sexual behavior (null), partial lordosis behavior, or complete (full) lordosis behavior in response to male attempts. Lordosis score and quotient were compared between groups afterward. The image was created with BioRender.com (license #DL273HIZWE).

To provide sexual experience, virgin proestrus females were mated with a sexually experienced male during the beginning of the dark period cycle and under dim red light in a silent room. The apparatus used was a clear acrylic open-field arena (70 x 70 x 35 cm) with shavings. The male was habituated to this cage for 10 minutes. After placing the female, the male was allowed to display up to 10 mounts with intromission attempts. At the same time, the characteristic sexually receptive lordosis behavior response was observed after each intromission. Only intromissions and no ejaculation occurred. The same procedure was repeated on the next proestrus phase of the cycle. Only females with a high lordosis behavior score and quotient (detailed below) were included in the following experimental step. I.e., females had to show an evident and complete (“full”) display of dorsiflexion with exposure of the genitals to facilitate intromission from 70 to 100% of the times the male attempted to copulate. This sexual female performance served as a control recording for testing the effects of microinjections into the MePD.

#### 2.3.3 Stereotaxic surgery and microinjection

Females were submitted to stereotaxic surgery on the day after the second mating session. They were deeply anesthetized (90 mg/kg ketamine and 10 mg/kg xylazine intraperitoneally) and implanted with a unilateral stainless steel 22-gauge guide cannulae directed to the right MePD, chosen in accordance to previous studies. Although both right and left MePD show plastic electrophysiological changes along the estrous cycle [8], females have a higher rate of synaptic vesicle release in the right MePD [52]. Moreover, when in proestrus, inhibitory synapses on dendritic shafts are higher than in the left MePD and higher than in the right MePD of males or diestrus and estrus females [52]. The right hemisphere also showed marked hormonal and neurochemical modulation of sexual behavior and cardiovascular reflex responses in males [10,53]. These findings addressed the right MePD *a priori* as a relevant site to test for its dynamic control of the behavioral display in females.

Therefore, the cannulae was placed 1 mm above the MePD at 2.9 mm posterior to the bregma suture, 4.1 mm lateral to the sagittal suture, and 5.1 mm below the dura mater, adapting coordinates from the atlas of Paxinos and Watson [48]. The guide cannulae were fixed to the skull with dental acrylic cement. Each rat received ketoprofen (3 mg/kg in 0.3 mL, i.p.), one injection per day for two subsequent days, and recovered 7 days before the following behavioral procedures. At 5 days post-surgery, vaginal cytology was re-evaluated. All females cycled normally.

In the next proestrus phase, females received a single microinjection into the MePD 3 h before the onset of the dark cycle, aiming to block the endogenous PRL effects during its first peak in circulation and coincidently with the preovulatory LH surge [20,21]. A tight-fitting 30-gauge infusion needle connected to a Hamilton microsyringe (USA) by a polyethylene tube was introduced into the stereotaxically implanted guide cannula tip. The microinjection needle extended 1 mm beyond the cannula tip. Microinjection was carried out for 60 s using a microinjection pump (Harvard Instruments, USA). The needle was left inside the cannula for an additional 60 s to prevent backflow of the microinjected substances.

One hour after the onset of the dark cycle, each female was placed together with a male, as mentioned above. All behavioral sessions were video recorded and further analyzed for quantifications of lordosis behavior. Sessions were limited to 10 intromissions by the male or to a maximum of 10 min recording time. The lordosis behavior display score was classified as “null”, “partial”, or “full” (depicted in Figure 1). I.e., in females with the highest sexual receptivity, all male attempts induced maximum lordosis (full display). In females with low sexual receptivity, either no significant increase in reflex dorsiflexion occurred during the copulatory attempt (null display), or the lordosis response was insufficient to allow penile intromission (partial display). This lordosis score was adapted from the proposed quantification by Hardy and DeBold [54]. Then, we also calculated the femalés lordosis quotient (LQ) using the formula [LQ = (number of “full” lordosis / 10 mounts) × 100], as performed in [55].

#### 2.3.4 Brain histology

The following day, females were sacrificed, as mentioned above, and the brains were removed from the skull and immersed in 10% formalin for one week. Using a vibratome (Leica, Germany), serial coronal sections (100 µm-thick) were obtained to determine the site of the implanted cannula and microinjection needle using a stereomicroscope (Olympus, Japan). As a rule, microinjections reached the medial border of the MePD, close to the optic tract and the adjacent sparse cells rim. Only animals with microinjections reaching the MePD [48] but showing no mechanical lesions or evident hemorrhages were included for data comparisons between experimental groups. Microinjections that were performed outside the borders of the MePD composed a heterogeneous “non-target” group whose results served as an additional approach to localize the effects tested in the area of interest (data not shown).

### 2.4 Statistical analysis

Data were analyzed using GraphPad Prism 8 or GraphPad InStat statistical software (USA). Mean values of gene expression results for each animal in each experimental group were calculated. Few values were considered outliers and excluded due to their evident deviation from the group data, also based on the ROUT test. The “n” (each one a pool of samples) for each tested gene ranges between 6 and 8 in the studied groups. These results were submitted to the Kolmogorov-Smirnov test for normality and Bartlett’s test for homogeneity of variances. Logarithm or square root transformations of the data were performed to meet the criteria for employing parametric tests when necessary. A two-way analysis of variance (ANOVA) test for repeated measures was used to compare separately each one of the gene expression by experimental group, hemisphere, and the interaction of these factors. Tukey’s *post hoc* test results are described separately for the right and the left hemispheres. We did not use the Bonferroni correction to avoid severely reducing the critical p-value. In addition, in cases where data transformation did not meet the conditions for parametric tests, the non-parametric Kruskal-Wallis (KW) test and Dunn’s *post hoc* test were used to analyze the experimental groupś data in the right and the left MePD. The KW test (corrected for ties) and Dunńs *post hoc* test were also used to compare both lordosis score and quotient values before and after microinjection between groups. The statistically significant level was set at p < 0.05 in all cases.

## 3. Results

### 3.1. Gene expression shows sex differences and hemispheric laterality effects in the MePD of males and cycling females

The RT-qPCR study was conducted to evaluate gene expression of *ERα, ERβ, GPER1, Kiss1, Kiss1R, PRGR, PRL, short* and *long PRLR, EGR1, JAK2, STAT5A,* and *STAT5B* in the right and left MePD of males and female in diestrus, proestrus, and estrus. Results (shown as the original, untransformed data) are presented in Figures 2-4.

**Figure 2.**
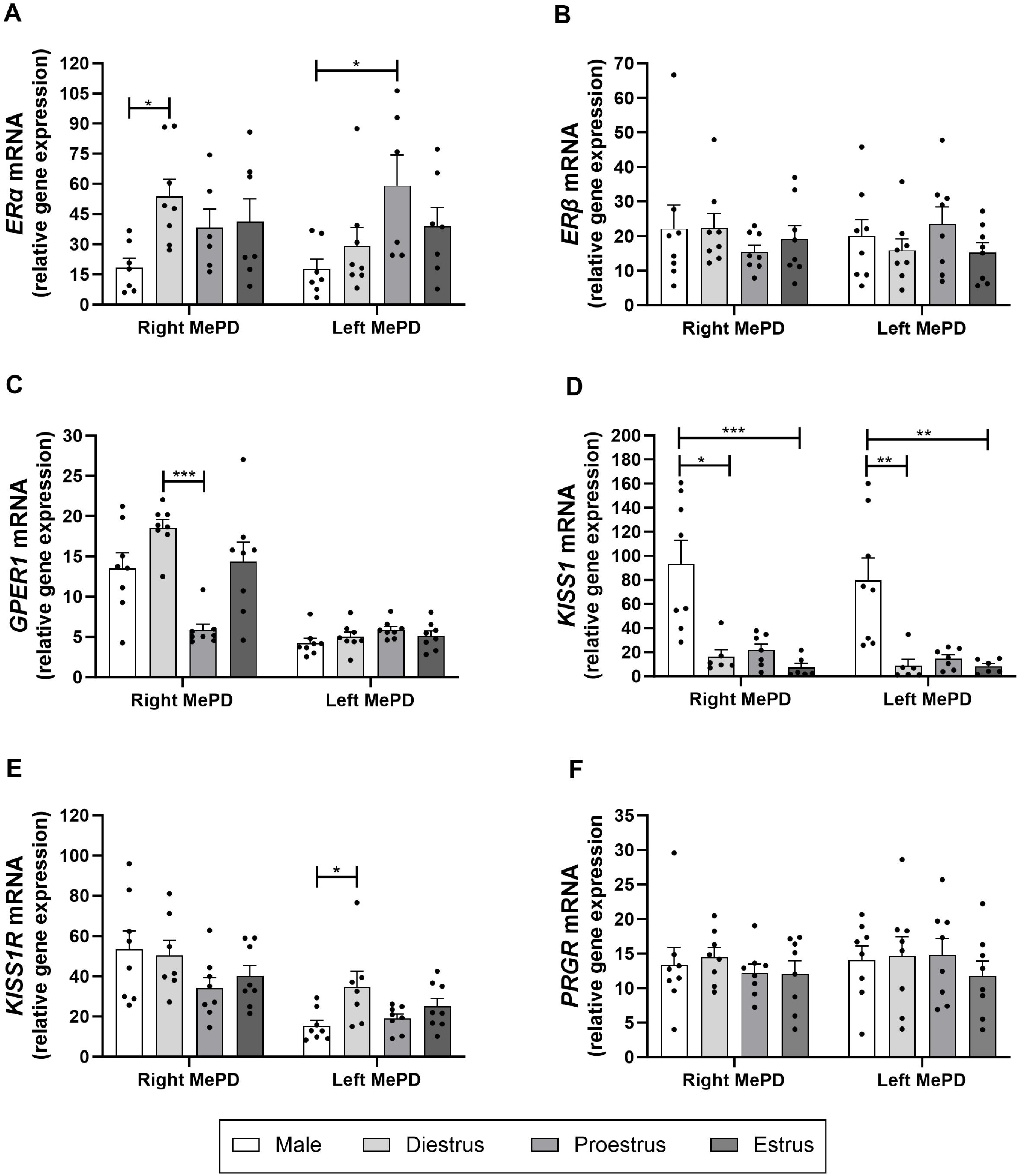
Relative gene expression of estrogen receptor alpha (*ERα;* **A**), estrogen receptor beta (*ERβ;* **B**), G protein-coupled estrogen receptor 1 (*GPER1;* **C**), kisspeptin (*Kiss1;* **D**), kisspeptin 1 receptor (*Kiss1R;* **E**), and progesterone receptor (*PRGR;* **F**) in the right posterodorsal medial amygdala (MePD) of male and female rats in different estrous cycle phases. (**A**) Diestrus and proestrus females show higher gene expression than male rats in the right and left MePD, respectively. (**B**) No statistically significant differences were found between groups and hemispheres (p > 0.05). (**C**) The right MePD of diestrus females has a higher gene expression than proestrus females. (**D**) Both the right and left MePD of male rats have a higher gene expression than the right MePD of diestrus and estrus females. (**E**) The left MePD of diestrus females has a higher gene expression than males. (**F**) No statistically significant differences were found between groups and hemispheres (p > 0.05). Data are presented as mean ± SEM. Points represent the data obtained for each sample in its respective experimental group, n= 6-8 per group. *\* p* < *0.05;* ** *p* < *0.01;* *** *p* < *0.001* for the comparisons indicated by brackets.

A statistically significant difference in the gene expression of *ERα* was found in experimental groups [F(3,24) = 5.230; p = 0.006] but not for hemispheres [F(1,24) = 0.054; p = 0.81] or in the interaction between these factors [F(3,24) = 1.598; p = 0.21]. The *post hoc* test showed lower values in the right MePD of males than diestrus females (p = 0.037). In the left MePD, males showed lower values than proestrus females (p = 0.019). Other group comparisons did not show significant differences (p > 0.12 in all cases; Figure 2A).

There were no statistically significant differences in the expression of *ERβ* between experimental groups [F(3,28) = 0.285; p = 0.83], hemispheres [F(1,28) = 0.13; p = 0.72], or in the interaction between these factors [F(3,28) = 1.02; p = 0.39; Figure 2B].

Gene expression of *GPER1* exhibited significant differences between experimental groups in the right MePD (KW = 15.724, p < 0.002). The *post hoc* test showed that values in diestrus females were higher than in proestrus (p < 0.001). Other comparisons did not show significant differences (p > 0.05 in all cases). The left MePD did not show a statistically significant difference between groups (KW = 7.114, p = 0.068; Figure 2C).

Gene expression of *Kiss1* showed significant differences between groups and hemispheres (KW = 17.089 and 17.312 for the right and left MePD, respectively; p < 0.001 in both cases). Comparing data by hemispheres, the *post hoc* test revealed that the values in the right MePD of males were higher than in diestrus (p < 0.05) and estrus (p < 0.001) but not in proestrus females (p > 0.05). In the left MePD, males also had higher values than diestrus and estrus females (p < 0.01 in both cases), but not when compared to proestrus ones (p > 0.05). Comparisons between females did not show significant differences in the MePD of both hemispheres (p > 0.05 in all cases; Figure 2D).

The *Kiss1R* gene expression did not show differences between groups in the right MePD (KW = 3.966, p = 0.265). In the left MePD, a statistically significant difference was found in the experimental groups (KW = 8.042, p = 0.045), and the post *hoc test* showed that diestrus females showed higher values than males (p < 0.05). Other comparisons did not reach the statistically significant level (p > 0.05 in all cases; Figure 2E).

The expression of *PRGR* was not different between experimental groups [F(3,28) = 0.586; p = 0.62], hemispheres [F(1,28) = 0.25; p = 0.61], or in the interaction between these factors [F(3,28) = 0.16; p = 0.91; Figure 2F].

Gene expression of *PRL* showed a statistically significant difference in experimental groups [F(3,21) = 5.939; p = 0.004], in the interaction of groups and hemispheres [F(3,21) = 3.886; p = 0.023], but not due to hemispheric laterality alone [F(1,21) = 0.004; p = 0.944]. The *post hoc* test showed that values in the left MePD of males were lower than in diestrus (p < 0.001), proestrus (p < 0.01), and estrus (p < 0.001) females. Other comparisons were not statistically significant (p > 0.05 in all cases; Figure 3A).

**Figure 3.**
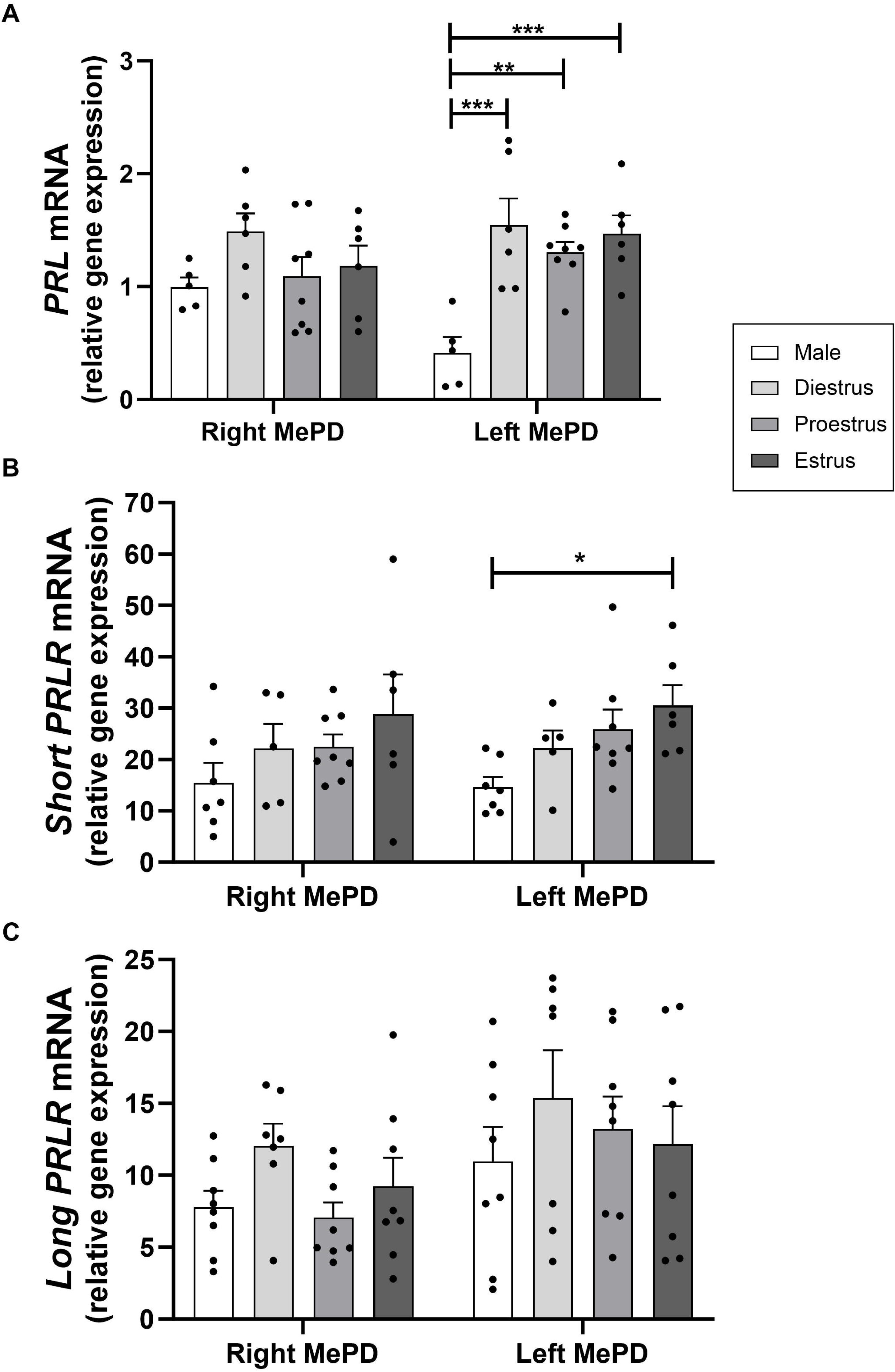
Relative gene expression of prolactin (*PRL;* **A**); prolactin receptor, short isoform (*Short PRLR;* **B**); and prolactin receptor, long isoform (*Long PRLR;* **C**) in the posterodorsal medial amygdala (MePD) of male and female rats in different estrous cycle phases. (**A**) Diestrus, proestrus, and estrus females have a higher gene expression in the left MePD compared to lower values found in males. (**B**) Estrous females has a higher gene expression than males in the left MePD. (**C**) No statistically significant differences were found between groups and hemispheres (p > 0.05). Data are presented as mean ± SEM. Points represent the data obtained for each sample in its respective experimental group, n= 5-8 per group. *\* p* < *0.05;* ** *p* < *0.01;* *** *p* < *0.001* for the comparisons indicated by brackets.

The expression of *Short PRLR* had a significant difference in experimental groups [F(3,22) = 4.758; p = 0.01] but not for hemispheric laterality [F(1,22) = 0.116; p = 0.73] or in the interaction of these factors [F(3,22) = 0.104; p = 0.95]. The *post hoc* tests revealed that values in the MePD of estrus females were higher than in males (p = 0.04). Other comparisons did not show statistically significant differences (p > 0.05 in all cases; Figure 3B).

The expression of *Long PRLR* was not different between experimental groups [F(3,27) = 1.024; p = 0.39], showed a statistically significant difference between hemispheres [F(1,27) = 11.95; p < 0.002], but the interaction of these factors was not statistically significant [F(3,27) = 0.467; p = 0.70]. The *post hoc* test did not find differences between the right and left MePD data (Figure 3C).

Likewise, the expression of *EGR1* was not different between experimental groups [F(3,28) = 0.96; p = 0.42], showed a statistically significant difference between hemispheres [F(1,28) = 6.178; p = 0.019], but the interaction of these factors was not statistically significant [F(3,28) = 1.056; p = 0.38]. The *post hoc* test could not identify the difference between the right and left MePD data (Figure 4A).

**Figure 4.**
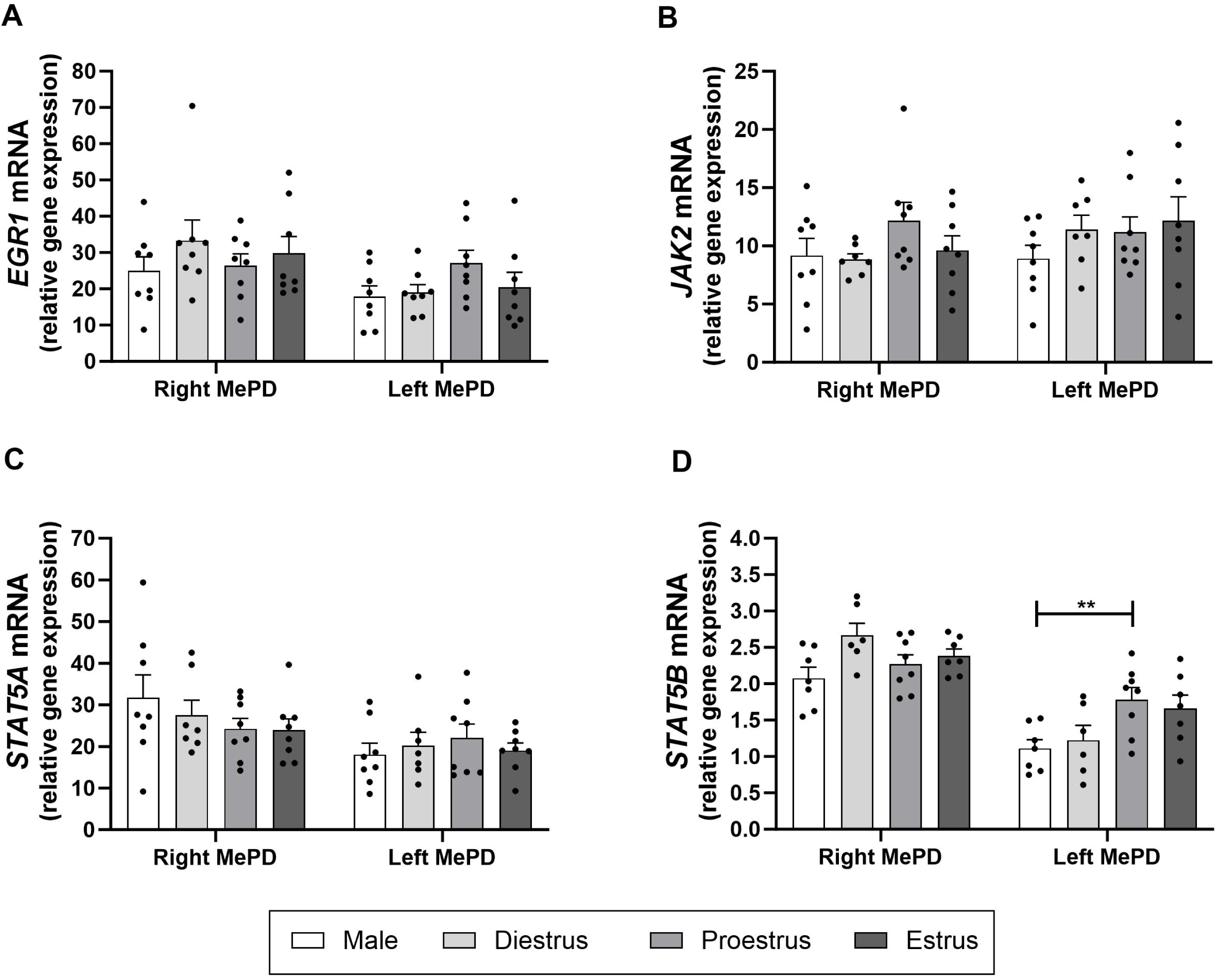
Relative gene expression of early growth response (*EGR1;* **A**), Janus kinase 2 (*JAK2;* **B**), signal transducer and activator of transcription 5A (*STAT5A*; **C**), and signal transducer and activator of transcription 5B (*STAT5B;* **D**) in the posterodorsal medial amygdala (MePD) of male and female rats in different estrous cycle phases. (**A, B, C**) No statistically significant differences were found between groups and hemispheres (p > 0.05). (**D**) Proestrus females have a higher gene expression than males in the left MePD. Data are presented as mean ± SEM. Points represent the data obtained for each sample in its respective experimental group, n= 6-8 per group. ** *p* < *0.01* for the comparisons indicated by brackets.

The expression of *JAK2* was not statistically different between groups [F(3,27) = 2.603; p = 0.07], hemispheres [F(1,27) = 0.636; p = 0.43], or in the interaction of these factors [F(3,27) = 0.599; p = 0.62; Figure 4B].

The expression of *STAT5A* was not different between experimental groups [F(3,27) = 0.484; p = 0.69], showed a statistically significant difference between hemispheres [F(1,27) = 7.115; p = 0.012], but the interaction of these factors was not statistically significant [F(3,27) = 0.911; p = 0.44]. The *post hoc* test did not identify the difference between the right and left MePD data (Figure 4C).

The expression of *STAT5B* showed significant differences between experimental groups [F(3,24) = 6.941, p = 0.001] and hemispheres [F(1,24) = 45.92, p < 0.001], but not in the interaction of these factors [F(3,24) = 2.266, p = 0.106]. The *post hoc* test showed higher values in proestrus females than males in the left MePD (p = 0.01). Other comparisons between groups were not significantly different but close to the statistical level of significance [males versus diestrus females in the right MePD (p = 0.058), and males versus proestrus and estrus females in the left MePD (p = 0.068 and 0.067, respectively; Figure 4D)].

In summary, the MePD expression of *ERα, GPER1, Kiss1, Kiss1R, PRL, Short PRLR,* and *STAT5B* differs in males and females. *ERα, Kiss1R, PRL, Short PRLR,* and *STAT5B* expression is higher in normally cycling females than males. *Kiss1* expression is higher in males than females, and *GPER1* is higher during diestrus than in proestrus.

### 3.2. Microinjection of Del1-9-G129R-hPRL into the MePD impairs the display of female sexual behavior

The right MePD modulates the sexual behavior display in rats, and the MePD has PRL-containing fibers and PRLR [34,35]. We tested the occurrence of estrogen-dependent lordosis behavior in proestrus females following the blockade of PRLR in the right MePD. Only results from rats whose microinjections reached this intended area without damaging it or the stria terminalis were selected for further study. Microinjection sites are shown in Figure 5.

**Figure 5.**
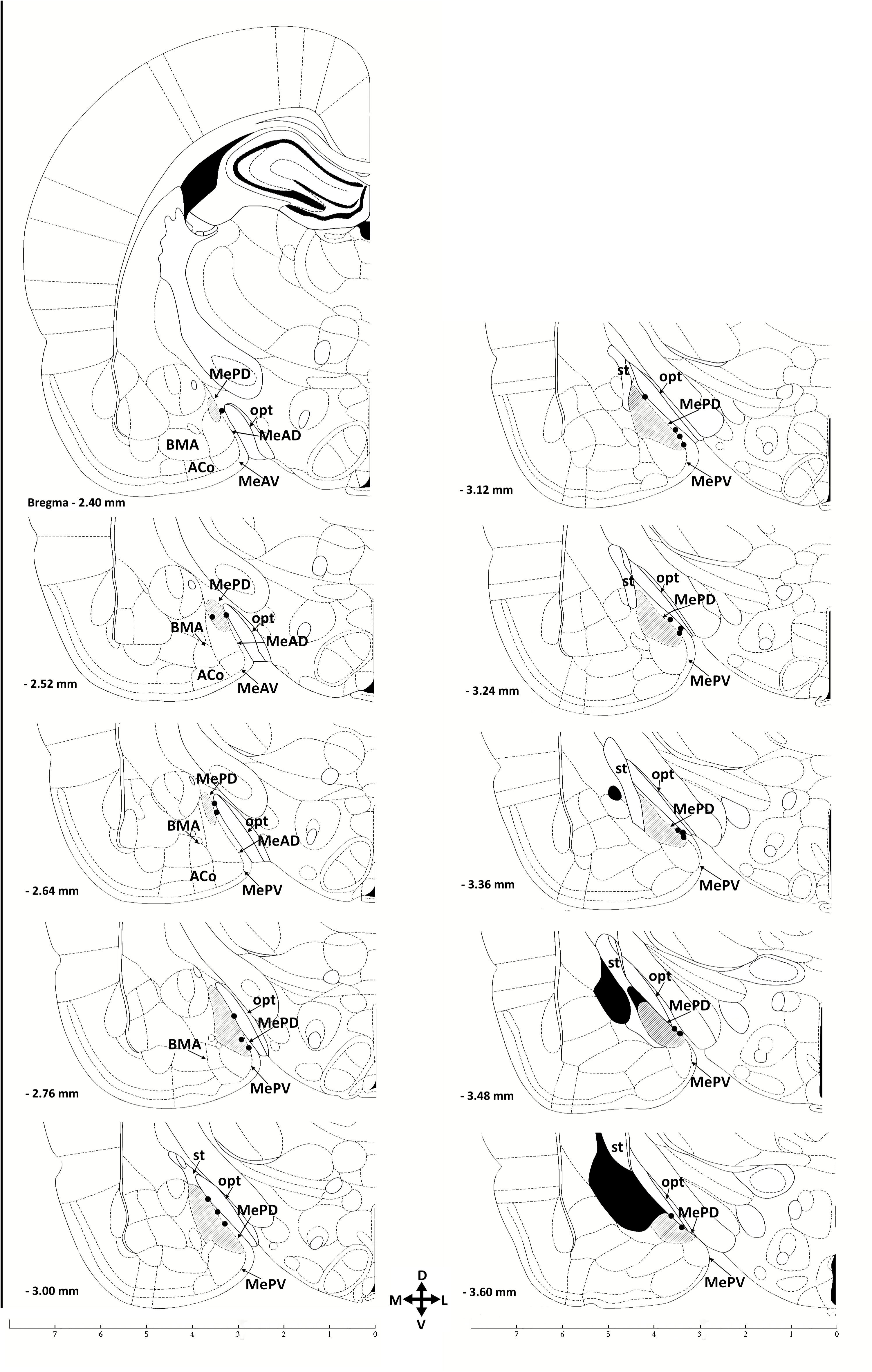
Schematic drawings of coronal sections showing the ventral forebrain of rats. The right MePD is highlighted in gray. Dots represent the approximate sites for the microinjections and spread of saline, 1 µM of Del1-9-G129R-hPRL, 10 µM of Del1-9-G129R-hPRL and co-injection of PRL 1 nM and Del1-9-G129R-hPRL 10 µM. No evidence of parenchymal lesions in the MePD were found in proestrus females included in the experimental groups. Coordinates are in millimeters posterior to the bregma (from 2.4 to 3.6 mm). ACo, anterior cortical amygdaloid nucleus; BMA, basomedial amygdaloid nucleus, anterior part; MeAD, anterodorsal medial amygdala; MeAV, anteroventral medial amygdala; MePD, posterodorsal medial amygdala; MePV, posteroventral part of the medial amygdala; opt, optic tract; st, stria terminalis. Adapted from the atlas of Paxinos and Watson [48].

The occurrence of lordosis behavior in the control recordings showed that females in all groups had similarly high rates of sexual receptivity in proestrus, clearly responding to the male attempts to copulate. There was a consistently high sexual receptivity in females of all groups, evidenced by a high lordosis behavior score (“full,” Figure 6A) and the calculated lordosis quotient (median values 90-100; Figure 6C). No statistically significant difference was found in the occurrence of complete lordosis behavior and lordosis quotient between groups in the control pre-microinjection recordings (KW = 6.134, p = 0.105 in both cases).

**Figure 6.**
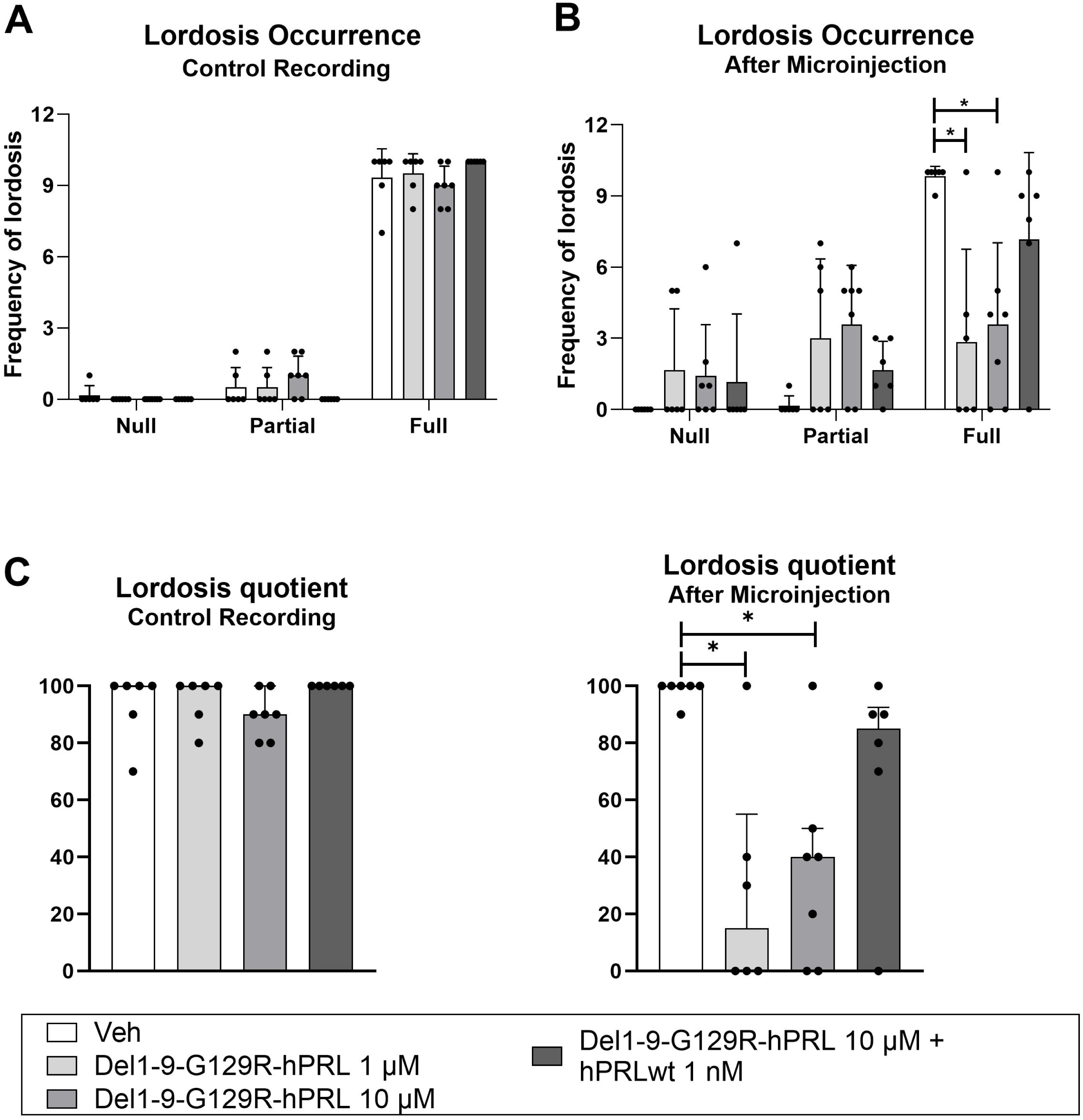
(**A**) Control recording of the sexual behavior (lordosis score) of proestrus female rats. The occurrence of lordosis behavior was classified as null, partial, or full display (depicted in Figure 1). All females showed higher levels of complete lordosis behavior. No statistically significant difference was found between females assigned to the different groups before the microinjection procedure (p > 0.05). (**B**) Lordosis score of proestrus female rats 3 hours after microinjection of saline, prolactin (PRL) receptor (PRLR) antagonist Del1-9-G129R-hPRL or the co-microinjection of PRL with PRLR antagonist into the right posterodorsal medial amygdala (MePD). Females that received 1μM and 10μM Del1-9-G129R-hPRL displayed a lower occurrence of full lordosis than the saline (Veh) group (p <0.05 in both cases). This inhibitory effect was reversed in the group that received a co-microinjection of PRL with the PRLR antagonist compared to Veh (p > 0.05). **(C)** Accordingly, the lordosis quotient was high during control recordings and showed a significant reduction in proestrus females microinjected with both doses of the PRLR antagonist (p < 0.05 in both cases), an effect reversed by the co-microinjection of PRL with Del1-9-G129R-hPRL when compared to the Veh group (p > 0.05). Data are presented as mean ± SEM in **A** and **B** and median and interquartile ranges in **C**; n 6-7 per group. Points represent data obtained from each female in each group. *\* p < 0.05* for the comparisons indicated by brackets.

Stereotaxic microinjections reached the MePD along its rostrocaudal axis (Figure 5). Statistically significant differences in lordosis score and lordosis quotient were found in proestrus females after microinjections (KW = 10.88, p = 0.0124 in both cases). The *post hoc* test demonstrated that the microinjections of 1 μM and 10 μM of Del1-9-G129R-hPRL in the MePD reduced the occurrence of full lordosis display and the lordosis quotient when compared to the control, Veh group (p < 0.05 in both cases). Indeed, these groups had reduced median values for lordosis quotient to 15-40 (Figure 6 B and C). This effect was reversed by the co-microinjection of PRL together with the PRLR antagonist (median value for lordosis quotient = 85, p > 0.05; Figure 6 B and C, respectively).

## 4. Discussion

The rat MePD has an expression of genes related to sex and estrous cycle stages and is a site for the modulatory role of PRL in the display of female sexual behavior. Our novel results demonstrate further MePD features with complex plasticity in the social behavior network, whose cells elaborate a dynamic “mosaic” of sex steroid actions and site-specific synaptic changes and plasticity [1,2,5–9,16,56]. These data are discussed and compared with relevant findings from other sex steroid-responsive brain areas as follows.

### 4.1. Sex steroid effects and gene expression in the MePD

Distinct gene expression and post-translation processing may involve different ERs in sexually dimorphic areas [4,56,57]. The induced expression of transcription factors and downstream signaling cascades contribute to a complex temporal organization of networks’ functioning. Differentially expressed mRNAs correlate better with their protein product than non-differentially expressed ones [58]. Then, the study of sex differences in mRNA levels provides an additional level of cellular functional organization for the MePD of males and cycling females [1,9,11,56].

Sex steroids control the development of neural circuits, regulate neuronal and glial shape and activity, and fine-adjust the responsiveness and the synaptic transmission of such networks [1,4,6,8,23,59–64]. From light and electron microscopy previous data, males and females in diestrus have a larger cell body volume [59], more proximal dendritic spines, and receive more excitatory synaptic inputs than proestrus and estrus females [5,8], while proestrus females show less dendritic spines (approximately 30% less than in diestrus) [5] and a higher density of somatic spines [7]. These cyclic changes altering the synaptic inputs to the MePD neurons and their output toward hypothalamic nuclei are related to the orchestrated sexual behavior display and ovulation in females [see additional data in 1,8].

Hemispheric laterality can also impose some differences in the MePD structure and function. The male MeA cells respond to both testosterone and estradiol [61,64]. Males have more neurons and astrocytes in the right MePD [65,66] and more complex astrocytes in the left MePD than females [66]. Rat MePD hemispheric asymmetries were also detected in electrophysiological recordings [more excitatory inputs in the left MePD of males; 8] and ultrastructural changes in cycling females [more inhibitory synapses on dendritic shafts in the right MePD of females in proestrus; 52]. Although there are some overlaps between hemispheric effects, it has been hypothesized that the left hemisphere has a greater steroid sensitivity, chemosensory processing, and control of LH secretion. In contrast, the right side would integrate the general, holistic aspects of the stimuli perceived [67]. Because the MePD sends only a few axons to each contralateral side, it is likely that the functional contributions of both right and left MePD are integrated into hypothalamic, brainstem, and cortical circuits onwards [17].

Multiple signaling mechanisms can coexist for the actions of gonadal hormones on target cells [e.g., ERα and ERβ involve KISS1 actions; 62]. Our data show that the MePD *ERα* and *Kiss1R*, as well as PRL, Short PRLR, and STAT5B expression, is higher in cycling females than males. *Kiss1* expression is higher in males than females, and *GPER1* is higher during diestrus than in proestrus. However, there is no simple scenario for the expression of genes in the MePD of adult males or in females along hours to a few days of specific estrous cycle phases. First, significant differences between males and cycling females involve the higher expression of *ERα* in females in diestrus (right MePD) and proestrus (left MePD). Estradiol levels stimulate this receptor, which has mutual interactions with ERβ and PRGR locally [4,63]. *ERβ* expression is sexually dimorphic in the MePD during the early postnatal hours, a finding that disappears by the fourth postnatal day because of a decrease in the expression of this receptor in females [62]. The difference in *ERα* expression remained constant over time [62], reinforcing the possibility that ERα causes main actions mediated by estradiol in the brain in adult rats and, then, ERα, but not ERβ, modulates female sexual behavior [15]. Here, *ERβ* expression in the MePD was not different in adult males and cycling females.

The *PRGR* expression was not sexually dimorphic in the MePD. Nevertheless, progesterone can exert local complex and dynamic structural actions together with estradiol across the estrous cycle or following ovariectomy and hormone replacement therapy [1,4,23,63]. Indeed, PRGR alters the density of dendritic spines in the female rat MePD differently under physiological conditions or in reorganized circuitries following castration [1,4,5,23].

The *GPER1* expression in the MePD is affected by sex, brain hemisphere, and estrous cycle phase. The right MePD of diestrus females showed higher expression than proestrus ones. GPER1 is related to rapid, non-genomic actions of estradiol and increases spine density in the hippocampal CA1 field of ovariectomized mice through phosphorylation of the actin polymerization regulator cofilin [68]. A reduced *GPER1* expression in proestrus is compatible with the reduction in the density of dendritic spines previously reported in the MePD of proestrus females compared to diestrus [1,5]. This finding might also correlate with the electrophysiological changes in proestrus [8] and, by reducing GABAergic output activity, the eventual disinhibition of female sexual behavior evidenced by the timely display of lordosis [1,5].

The expression of *Kiss1* develops along the ontogeny and is sexually dimorphic in the MeA of rats and mice [31,32]. *Kiss1* levels in the MeA are upregulated by gonadal hormones, and mice lacking kisspeptin or KISS1R show impairments in reproduction and fertility [29]. In the male MeA, androgen and estrogen converted by aromatase [29,70,71] might modulate *Kiss1* expression and the display of male sexual behavior [1,16,52]. Here, the *Kiss1* expression in male rats is higher than in diestrus and estrus females in both hemispheres. *Kiss1R* expression is higher in diestrus females than males in the left MePD. These data indicate a sex difference in both *Kiss*1 and *Kiss1R* expression in the MePD, which is sensible to estradiol and progesterone levels but are not similarly modulated by them, a finding also dependent on the brain hemisphere [expanding the data in 32,69].

Estradiol enhances GnRH-induced EGR-1 mRNA and protein expression in mouse gonadotrope cell line culture [72], and both ERα and ERβ regulate EGR1 mRNA in the ovariectomized rat pituitary gland [73]. Although involved in the molecular mechanism of transcription and the hypothalamic-pituitary-ovary axis activity, *EGR1* expression showed no significant differences in sex or estrous cycle phase in the MePD. This region-specific effect is interesting because *EGR1* expression in the anterior pituitary of female rats was higher during proestrus than in diestrus [73] and temporally associated with an increase in pituitary PRL synthesis [74] and the peak of circulating PRL during proestrus [20]. It is interesting that, although the estrous cycle alone cannot alter *EGR1* expression in the MePD, selective EGR-1 responses are induced in this area of proestrus females by the neural processing of vaginocervical stimulation (VCS) during mating [75]. This latter finding indicates that EGR-1 activity is stimuli-specific in the female MePD and depends on the associated occurrence of proestrus, full lordosis display, and sensory input related to copulation.

### 4.2. Integrated PRL actions in the MePD

Previous data indicated that PRL is related to maternal aggression and stress responses acting in the central nucleus of the amygdala of mice and rats [74]. We tested an additional amygdaloid area and functional role for PRL. The present findings indicate that the plastic, sex steroid-responsive MePD is also sensitive to PRL to modulate sexual receptivity in proestrus females.

We studied the blockage of PRLR in the female MePD in parallel to the gene expression experiments. The right MePD was chosen *a priori* for microinjection for its marked changes in synaptic processing during proestrus and its role in sexual behavior display, as mentioned above. Indeed, estrogen has multiple modulatory effects in sensitive brain areas of normally cycling females [76]. Tissue-specific and site-specific PRL expression and actions also occur [45,74,77,78]. Notably, the female MeA contains a high density of both ER and PRLR, and local PRL-induced pSTAT5-immunoreactive neurons coexpress ERα [77]. Interestingly, males had a distinct, lower *PRL* expression in the left MePD than females. In contrast, females maintained local values stable along the estrous phases in both hemispheres. Comparing the two isoforms of the PRLR, the *Short PRLR* expression showed sex differences (higher in estrus) but not the *Long PRLR* one. Since circulating PRL increases along with proestrus, higher levels would reach the female MePD and add to the “basal” production of local PRL. Afterward, different post-transcriptional mechanisms would integrate the actions of sex hormones, PRL, and PRLR to initiate intracellular signaling sequences, such as the ERα actions on STAT5 transcriptional activity induced by PRL-PRLR-JAK2 cascade.

Again, site-specific features are evident because while estradiol alters *Long PRLR* expression in the hypothalamus [41 and references therein] and PRL synthesis is modulated by ERβ, but not by ERα, in the paraventricular and supraoptic hypothalamic nuclei of female rats [78], neither *ERβ* nor *Long PRLR* expression showed differences between experimental groups in the MePD. Integrated neuronal networks’ PRL actions can be even more complex, as seen for the maternal aggression display [79; see also 5 for further discussion], or in another functional scenario that occurs in male rats, for the MeA evokes a marked increase in serum PRL following fearful stimuli and stressful conditioning [80].

On the other hand, the expression of both *JAK2* and *STAT5A* was also not different between groups, but the expression of the *STAT5B* gene was higher in the left MePD of proestrus females than in males. This finding aligns with the abovementioned *PRL* and *Short PRLR* expression differences between females and males, pointing to STAT5B as a candidate to intracellularly modulate the local effects of PRL during its peaks in circulation [see also 25]. STAT5B serves the PRL-mediated negative feedback in tuberoinfundibular dopamine neurons [81] and is the mediator (not STAT5A) of PRL actions in the hypothalamus [25,82]. These findings also address the participation of the left MePD projections in the central pathways for neuroendocrine release control while the right MePD would display a modulatory effect of PRL on the occurrence of female sexual behavior, as tested here.

We studied the effect of a PRLR antagonist on the sexually receptive behavior displayed in response to the malés attempts to copulate. This was relevant because PRL levels increase around the time of ovulation, the proestrus surge of PRL enhances female sexual receptivity [46], and bilateral MePD excitotoxic lesions abolish the preovulatory PRL surge in the afternoon of proestrus [83]. Here, the typically high lordosis score and quotient were significantly reduced following the MePD microinjection of the PRLR antagonist Del1-9-G129R-hPRL (1µM and 10µM) in the afternoon of proestrus. This behavioral impairment was reversed by the co-microinjection of the receptor antagonist combined with PRL, assessing the PRLR dependence of this effect [47]. That is a novel role for the PRL action in the MePD and its participation in the central network that dynamically modulates the reproductive behavior of females. They can also be associated with results showing that, in proestrus females, the MePD connects with medullar and forebrain networks for transducing VCS during mating into PRL release required for the maintenance of pregnancy or pseudopregnancy [84]. These data add to the sex differences in postpubertal functioning of the MePD [85], and address further approaches to investigate whether PRL effects in the MePD are synchronized, additive, or redundant for the temporal organization of the timely female sexual display and neuroendocrine control of ovulation during proestrus.

## 5. Conclusion

We studied the gene expression of *ERα, ERβ, GPER1, Kiss1, Kiss1R, PRGR, PRL, Short PRLR, Long PRLR, EGR1, JAK2, STAT5A,* and *STAT5B* in the MePD of both hemispheres in adult males and females in diestrus, proestrus, and estrus. The expression of *ERα, GPER1, Kiss1, Kiss1R, PRL, Short PRLR*, and *STAT5B* showed sex differences, suggesting a different role for these genes in both sexes and a rapid, dynamic switch across the estrous cycle phases. Likewise, in proestrus females, PRL action in the MePD is needed for the display of sexual behavior when high levels of ovarian steroids are in circulation. These data add to the plastic morphological and electrophysiological fine-tuning of the MePD related to reproduction within the social neural network of adult rats.

## Data availability statement

All original data are published here. Raw data will be available under request.

## Ethics statement

The Ethics Committee on the Use of Animals at the Federal University of Health Sciences of Porto Alegre (Brazil) reviewed and approved this study (#616/19 and 641/19). No generative artificial intelligence was used to write this manuscript.

## Author contributions

VG, AARF, LB: study conceptualization and design; AARF, LB*: in vivo* experimental procedures; LB: *ex vivo* tissue and sample analysis; LB, ACM, MG: molecular procedures; AARF, MG, LB: statistical analysis; VG, AARF, MG: critical revision of the manuscript. All authors have written, read, edited, and approved the final version of the manuscript.

## Acknowledgments

**The authors thank Mrs. Carmen Lúcia Andrades, Alexandra Guimarães, and Milena Silveira for technical support, Dr. Clarissa Hollenbach (UFCSPA, Brazil) for skilled animal care,** and Lucas Bühler for his help with the figures.

## Abbreviations

Del1-9-G129R-hPRL: prolactin receptor antagonist
EGR1: Early Growth Response Factor 1
ERα: α-estradiol receptor
ERβ: β-estradiol receptor
FSH: follicle stimulating hormone
GnRH: gonadotropin-releasing hormone
GPER1: G protein-coupled estrogen receptor
i.p.: intraperitoneal
JAK2: Janus kinase 2
KISS1: kisspeptin 1
KISS1R: kisspeptin 1 receptor
LH: luteinizing hormone
MAP-kinase: mitogen-activated protein kinase
MePD: posterodorsal subdivision of the medial amygdaloid nucleus
PRGR: progesterone receptor
PRL: prolactin
PRLR: prolactin receptor
RT-qPCR: real-time quantitative polymerase chain reaction
STAT5: signal transducer and activator of transcription 5
TIDA, VCS: vaginocervical stimulation
Veh: Vehicle

